# Predicting plant-related and leaf-related parameters of winter wheat using vegetation index: Should phenological correction be applied across growth stages?

**DOI:** 10.1101/2022.03.07.483295

**Authors:** Yu Zhao, Yang Meng, Haikuan Feng, Shaoyu Han, Guijun Yang, Zhenhai Li

## Abstract

Most existing plant-related and leaf-related parameters models of winter wheat vary across growing seasons, but it is an open question whether a unified statistical model can be developed to predict plant-related and leaf-related parameters using VI across multiple growing seasons, or whether the phenological correction is necessary for each parameter across multiple growing seasons. To explore this question, we measured two plant-related parameters and four leaf-related parameters over five growth stages during the 2017–2021 growing seasons. A hierarchical linear model (HLM) automatically adapts the relationship between VIs and their corresponding parameter across growing seasons and assesses the contribution of phenological variables by applying a sensitivity analysis. The estimates of VI–plant-related parameters [aboveground dry biomass (AGB) and plant nitrogen concentration (PNC)] were scattered over a given growing season, unlike the relationship between VI–leaf-related parameters [leaf dry biomass (LGB), leaf nitrogen concentration (LNC), leaf area index (LAI) and soil and plant analysis development (SPAD)]. In contrast, the AGB, PNC, LGB, LNC, LAI, and SPAD HLM models are stable and can be popularized across growing seasons, with the determination coefficient *R*^2^ ranging from 0.84 to 0.86, 0.79 to 0.87, 0.70 to 0.71, 0.68 to 0.86, 0.75 to 0.81, and 0.68 to 0.70, respectively, and the root mean square error ranging from 0.13 to 0.50 t/ha, 0.01 to 0.07%, 0.01 to 0.03 t/ha, 0.02 to 0.03%, 0.02 to 0.07, and 0.47 to 0.69, respectively. The sensitivity index of the phenological information in the AGB and PNC models was 0.56–0.78 and 0.66–0.72, respectively, whereas that in the LGB, LNC, LAI, and SPAD models was 0.01–0.06, 0.01–0.10, 0.02–0.06, and 0.00–0.01, respectively. Although phenological effects have little effect on leaf-related indicators, HLM has a strong potential for application to other crops and regions.

## 1 Introduction

Crop-growth monitoring provides a basis for real-time knowledge of seedling conditions, farmland management, and early-yield estimation and has become an important part of precision agriculture (Guo et al., 2019). Advances in remote-sensing technology have led to the development of various statistical and physical methods to retrieve agronomic parameters (APs) (Eitel et al., 2014; Nguy-Robertson et al., 2014; Berger et al., 2020). However, the use of similar spectra of various objects over multiple growing seasons remains an open issue in the field of agronomic-parameter monitoring.

Tracking variations in plant- and leaf-related parameters over a growing season is critical to understanding the variations in the spectra of growing vegetation. Many APs reveal allometric accumulation or reduction across growth stages due to material accumulation and transportation (Zhao et al., 2021), whereas the vegetation index (VI) varies quadratically with the growth period (Lee et al., 2016; Luo et al., 2020). One set of VIs may correspond to multiple APs, such as aboveground dry biomass (AGB) or plant nitrogen concentration (PNC) (Yue et al., 2017; Zheng et al., 2018). Most of the literature concords that, although VIs may correlate well with APs over a certain growth stage, they do not transfer to other times or locations (Stroppiana et al., 2009; Yu et al., 2013; Zhou et al., 2018; Feng et al., 2019; Li et al., 2022). Substantially the piecewise method provides more accurate and useful phenological information (Lee et al., 2016; Din et al., 2017).

Unfortunately, the discontinuity of the piecewise method can lead to uncertainty in the prediction model for uninvestigated periods. When dealing with multiple growth stages, APs based on VIs are difficult to estimate for three reasons: (i) the winter wheat canopy structure and background information in the field observation change constantly, which causes the sensitivity of VIs to APs to depend on the growth stage for which the VI is constructed, (ii) optical bands and VIs saturate at medium-to-high canopy cover, and (iii) optical VIs are not sensitive to obscured information, such as information from the lower layers of vegetation close to ground and stems.

Numerous anti-saturation VIs and VIs that eliminate soil background have been constructed, such as the optimized nonlinear VI (Feng et al., 2019) and the optimized soil-adjust VI (Rondeaux et al., 1996). Several remarkable improvements have been made for estimating AGB, PNC, and leaf area index (LAI) from the image textural characteristics, vertical distribution rule, and plant height, or by combining synthetic aperture radar and spectral information, which corrects the phenological difference in the prediction (Nguyen and Lee, 2006; Tian et al., 2014; Zheng et al., 2018). The uncertainties of the radiation transfer model were reduced by varying the model parameters over the growth stages (Li et al., 2018). However, the robustness of these new techniques and methods is limited by cost, flexibility, data-processing difficulties, and image quality.

In addition, deep-learning algorithms are widely used to predict APs with excellent results. A well-trained network depends on a wealth of training datasets that are expensive to collect, and their generality for image quality has not been thoroughly tested. We know of only two studies that examined how to automatically adapt the relationship between VIs and APs across growing seasons for plant-related parameters, including AGB and plant area index (Li et al., 2022). No study has yet focused on whether a unified statistical model can be developed to predict plant-related and leaf-related parameters using VI, or whether a phenological correction is required for each parameter across multiple growing seasons.

To respond to this need, we focus herein on estimating plant-related parameters (AGB and PNC) and leaf-related parameters [leaf dry biomass (LGB), leaf nitrogen concentration (LNC), leaf area index (LAI), and soil and plant analysis development (SPAD)] for winter wheat throughout the entire growing season by using five well-known VIs. The specific research objectives are (1) to investigate the relationship between VI and the parameters over individual growth stages or entire growth stages; (2) to construct a VI-based adaptation approach to predict plant- and leaf-related parameters and examine the accuracy of such predictions; and (3) to evaluate how phenological information and VIs contribute to different AP models.

## 2 Materials and methods

### 2.1. Study sites and experimental design

The experiments were conducted over a four-year period at Xiaotangshan National Precision Agriculture Research Center (40.17°N, 116.43°E, average altitude: 36m) near Beijing (Figure 1). Experiments (Exp) were conducted over four consecutive seasons from 2017 to 2021 at the Xiaotangshan National Precision Agriculture Research Center in the suburb of Beijing, China (40.17°N, 116.43°E) (Figure 1c, Reg. 2). Table 1 details the experimental design.

**Figure 1.**
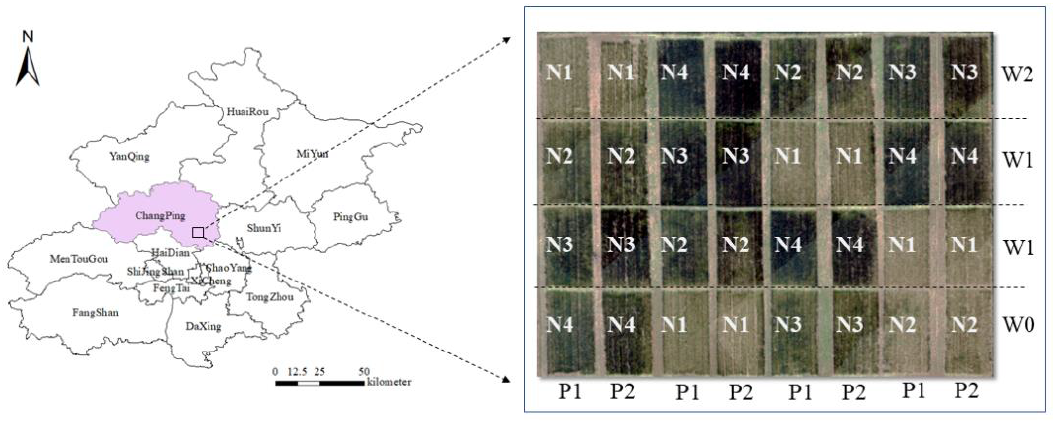
Location of Xiaotangshan Experimental Station. Experimental designs P, N, and W represent different varieties, nitrogen rates, and irrigation rates over different growing seasons.

**Table 1.**
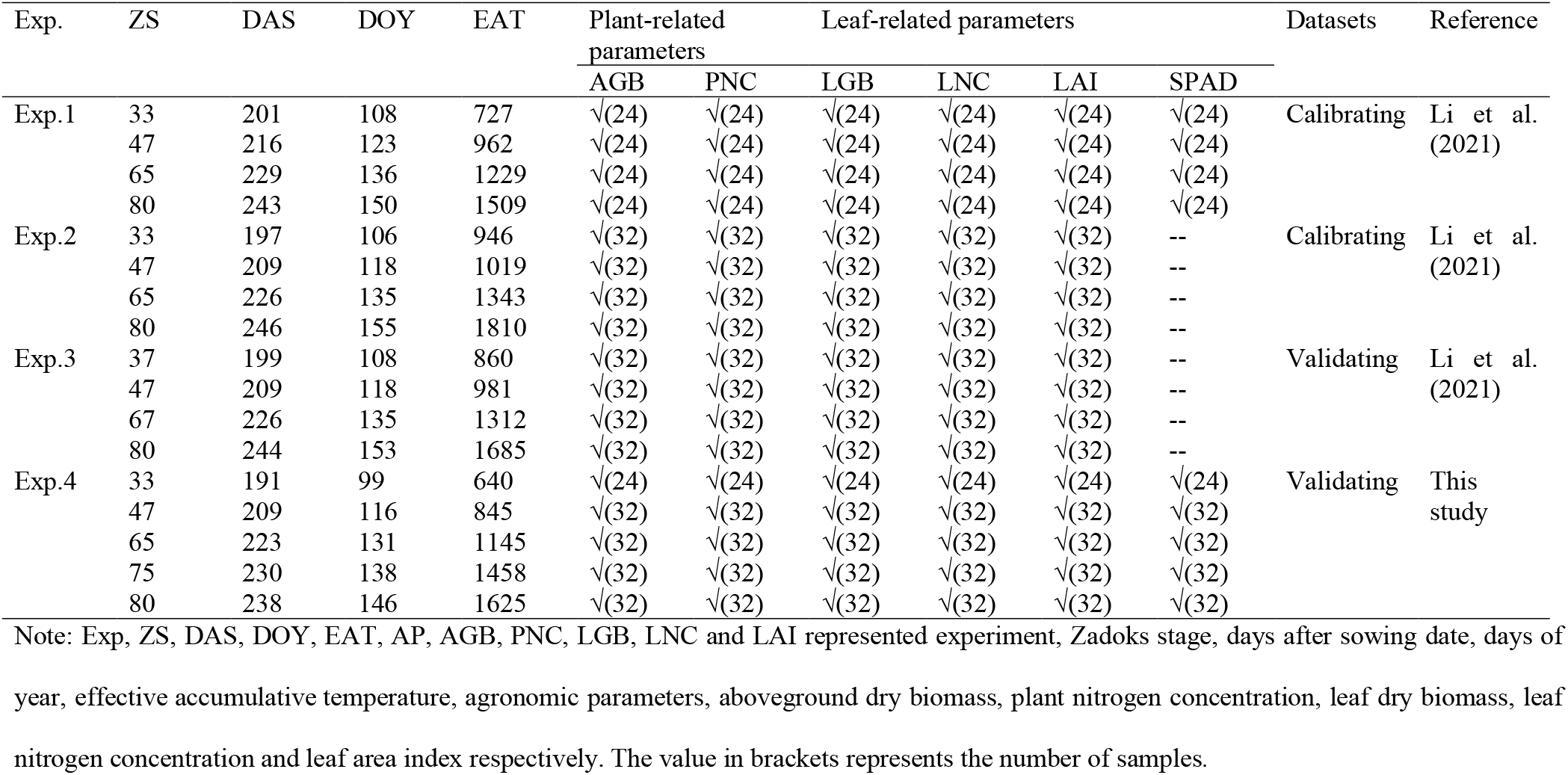
Results of winter-wheat field measurements made over different growing seasons.

Three experiments, 2017–2018 (Exp. 1), 2018–2019 (Exp. 2), and 2019–2020 (Exp. 3), used completely randomized designs (Li et al., 2021), whereas the growing season 2020–2021 (Exp. 4) was designed as a three-way factorial arrangement of cultivars (P1: Jinghua 11 and P2: Zhongmai 1062), nitrogen (N1: 0 kg N ha^−1^; N2: 90 kg N/ha; N3: 180 kg N/ha, and N4: 270 kg N/ha) and irrigation (W0: 25 mm, W1: 148 mm, and W2: 241 mm) treatments in a split-plot design (Figure 1). Different years, sample dates, and sample numbers were used for winter wheat and spring wheat (Table 1).

### 2.2 Data sources

#### Plant measurements

Twenty tillers of winter wheat plants per experimental plot were randomly sampled over different growth stages to determine plant-related parameters (AGB and PNC) and leaf-related parameters (LGB, LNC, LAI, and SPAD). The SPAD is the mean value obtained from measuring the leaf value of the 20-stem winter wheat plant by using a SPAD-502 instrument (Minolta Camera Co. Ltd., Japan). Table 1 gives the number of samples collected in each experiment. The measurement methods and equations for calculating AGB, LGB, PNC, LNC, and LAI are the same as used by Zhao et al. (2020).

#### Field canopy hyperspectral data

The canopy reflectance spectra were collected from 2017 to 2021 at the Xiaotangshan experimental plots by using a Field Spec Pro FR2500 (Analytical Spectral Devices, Boulder, CO, USA). Measurements were made during cloud-free periods between 10:00 a.m. and 2:00 p.m. to minimize variations in illumination conditions. Before measuring the canopy reflectance, a white Spectrolon reflectance panel (Labsphere, North Sutton, NH, USA) with diffuse reflectance properties was used to calibrate the spectral reflectance. The instrument was held 1.0 m above the canopy, and the view nadir and azimuth angles were both zero. For each plot, hyperspectral data were collected from one representative 1.0 m×1.0 m sub-plot. To reduce the effects of field conditions, ten scans were acquired and averaged to represent the canopy reflectance of each plot.

### 2.3 Estimation method and procedure

#### Spectral vegetation indices

We selected five VIs as good candidates for evaluating plant-related parameters (AGB and PNC) and leaf-related parameters (LGB, LNC, LAI, and SPAD) (Table 2) and analyzed the correlations between different APs and each selected VI. Finally, the optimal VIs were used to estimate how the phenology affects the hyperspectral inversion of APs.

**Table 2.** Summary of spectral vegetation indices used in this study.

| Index | Formula | Reference |
| --- | --- | --- |
| Normalized difference vegetation index (NDVI) | $NDVI = (R_{800} - R_{670}) / (R_{800} + R_{670})$ | Rouse et al., 1974 |
| Optimized soil-adjusted vegetation indices (OSAVI) | $OSAVI = (1 + L) * (R_{800} - R_{670}) / (R_{800} + R_{670} + L)$ | Rondeaux et al., 1996 |
| Green normalized difference vegetation index (GNDVI) | $GNDVI = (R_{750} - R_{550}) / (R_{750} + R_{550})$ | Gitelson., 1996 |
| Enhanced vegetation index 2 (EVI2) | $EVI2 = 2.5 * (R_{800} - R_{670}) / (R_{800} + 2.4 * R_{670} + 1)$ | Jiang et al., 2008 |
| Photochemical reflectance index (PRI) | $PRI = (R_{531} - R_{570}) / (R_{531} + R_{570})$ | Gamon et al., 1992 |

Note: $R_{531}$ , $R_{550}$ , $R_{571}$ , $R_{670}$ , $R_{750}$ and $R_{800}$ represent the reflectance at 531 nm, 550 nm, 571 nm, 670 nm, 750 nm and 800 nm, respectively. L is the soil adjustment coefficient, which is 0.16 in this paper.

#### A Method based on vegetation indexes to adapt the phenology for predicting winter-wheat parameters

Given that the statistical model parameters for plant- and leaf-related parameters over the various phenological periods are different (Table 4), we bridged the VIs and the phenology and APs (AGB, PNC, LGB, LNC, LAI, and SPAD) to automatically adjust the VI-AP relationships as a function of phenology. The hierarchical linear model (HLM) was introduced to analyze the phenological correction (Lininger et al., 2015). At level 1, the APs are linear in the phenological stage. The function is characterized by a set of parameters, and the HLM is given by

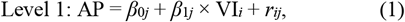

where AP, *β*_0*j*_, and *β*_1*j*_ are the APs (including AGB, PNC, LGB, LNC, LAI, and SPAD), the intercept, and the slope of the linear model, and *r_ij_* is the random error.

At level 2, these parameters vary across phenological stages, and the parameters of the first layer are automatically corrected by the phenological data of the second layer:

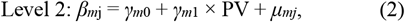

where *β_mj_* corresponds to *β*_0_ and *β*_1_ from the HLM model, respectively, *γ*_*m*0_ is the intercept, *γ*_*m*1_ is the slope of each PV and *μ_mj_* is the random error of Level 2 of the HLM. PV is the set of phenological variables [Zadoks stage (ZS), days after sowing (DAS), day of year (DOY), and effective accumulative temperature (EAT)], which are used to quantify the phenological development process of winter wheat. The ZS period is the field survey income according to the definition of Zadoks et al. (1974). The EAT is calculated as follows:

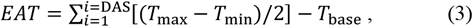

where *T*_max_ and *T*_min_ are the maximum and minimum daily temperatures, respectively. Moreover, *T*_max_ = *T*_base_ if *T*_max_ < *T*_base_. For winter wheat, *T*_base_ = 0 °C (Acevedo et al., 2009).

#### Linear models

To assess how phenology affects estimates of the plant- and leaf-related parameters, we constructed and compared a multiple linear regression (MLR) of a piecewise linear model based on different growth stages.

#### Extended Fourier amplitude sensitivity test and sensitivity analysis

To analyze the contribution of VIs and of the phenological information of the HLM, we applied an extended Fourier amplitude sensitivity test, which combines the Fourier amplitude sensitivity test with the Sobol algorithm to determine the sensitivity index to quantify how input variables depend on output variables (Saltelli et al., 1999). This procedure involves four steps: (1) the independent variables (VI and PV) are defined, and their ranges are from the maximum to the minimum values of the measured samples. (2) The set of independent variables are generated by Monte Carlo sampling of distributions, generating a total of 100 000 datasets. (3) The predicted AP for each sample set of independent variables were calculated by invoking the HLM model in Matlab 2016 (Mathworks, Inc., Natick, MA, USA) programming. (4) The sensitivity index of each independent input variable to the APs is computed by using Simlab (ver 2.2.1). The total variation is

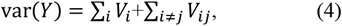

where *V_i_* is the variability associated with each independent variable of the HLM model, and *V_ij_* is the variability associated with the interaction between each independent variable of the HLM model. The sensitivity index is derived from the decomposition of Eq. (4) by dividing the importance measures by var(Y):

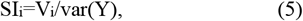

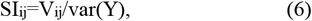

where S_i_ is also called the sensitivity index, and *S_ij_* is the interactive sensitivity index between the VI and PV. The sensitivity index of each independent input variable is computed by using Simlab (ver 2.2.1).

#### Model evaluation

This paper analyses the correlation between the VI and various APs over different growth stages based on a Pearson correlation analysis. To test whether a significant difference exists in the regression relationships under different phenological conditions, we examine the difference in regression coefficients by using the F-test. We evaluate the model performance by using the determination coefficient *R*^2^ and the root mean squared error (RMSE). These are calculated as follows:

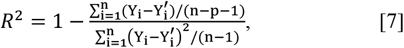

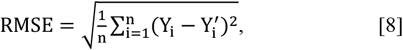

where *n*, *Y*′_*i*_, *Y_i_*, and *p* are the number of samples, predicted value, measured value, and the number of independent variables, respectively.

## Results

### 3.1 Correlation between agronomic parameters and vegetation indexes

Five commonly used VIs were selected to establish relationships with plant-related parameters (AGB and PNC) and leaf-related parameters (LGB, LNC, LAI, and SPAD) by using data from the 2017 to 2021 growing seasons of winter wheat (Table 3). All selected VIs correlate weakly with AGB, with the OSAVI having the strongest correlation (*r* = 0.18), followed by the EVI2 (*r* = 0.11) and the PRI (*r* = −0.10). The VI with the highest correlation with PNC is the GNDVI (r = 0.50), followed by the normalized difference vegetation index (NDVI; *r* = 0.48), the PRI (*r* = 0.48), the EVI2 (*r* = 0.46), and the OSAVI (*r* = 0.41). VIs correlate significantly with LGB, LNC, LAI, and SPAD, with the strongest correlation being OSAVI (*r* = 0.77), PRI (*r* = 0.78), OSAVI (*r* = 0.80), and OSAVI (*r* = 0.81), respectively. Therefore, the OSAVI, OSAVI, GNDVI, PRI, OSAVI, and OSAVI are used for subsequent analysis of AGB, LGB, PNC, LNC, LAI, and SPAD.

**Table 3.** Correlations between agronomic parameters and VIs.

| VI | Plant-related parameters |  | Leaf-related parameters |  |  |  |
| --- | --- | --- | --- | --- | --- | --- |
|  | AGB | PNC | LGB | LNC | LAI | SPAD |
| NDVI | 0.08 | 0.48 | 0.75 | 0.73 | 0.76 | 0.78 |
| OSAVI | 0.18 | 0.41 | 0.77 | 0.74 | 0.80 | 0.81 |
| GNDVI | 0.03 | 0.50 | 0.74 | 0.75 | 0.75 | 0.72 |
| EVI2 | 0.11 | 0.46 | 0.75 | 0.69 | 0.80 | 0.76 |
| PRI | -0.10 | 0.48 | 0.75 | 0.78 | 0.77 | 0.75 |

### 3.2 The performance of vegetation indexes and agronomic parameters over different phenological stages

Experimental data from seven growth stages (ZS 33 to ZS 80) were collected, and the range of AGB, LGB, PNC, LNC, LAI, and SPAD of the entire dataset was 0.52– 22.62 t/ha, 0–3.82 t/ha, 0.86%–3.95%, 1.05%–5.07%, 0%–7.34, and 3.7–56.77, respectively (Figure 2). All vegetation indexes tested herein are closely connected to plant-related parameters (AGB and PNC) and leaf-related parameters (LGB, LNC, LAI, and SPAD) with correlation coefficients *r* in each ZS exceeding 0.41 (Table 4). No significant difference appears in the correlation of AGB, PNC, LGB, LNC, LAI, and SPAD with selected VIs, ranging from 0.61 to 0.89, 0.41 to 0.90, 0.53 to 0.88, 0.55 to 0.89, 0.75 to 0.87, and 0.64 to 0.92, respectively.

**Figure 2.**
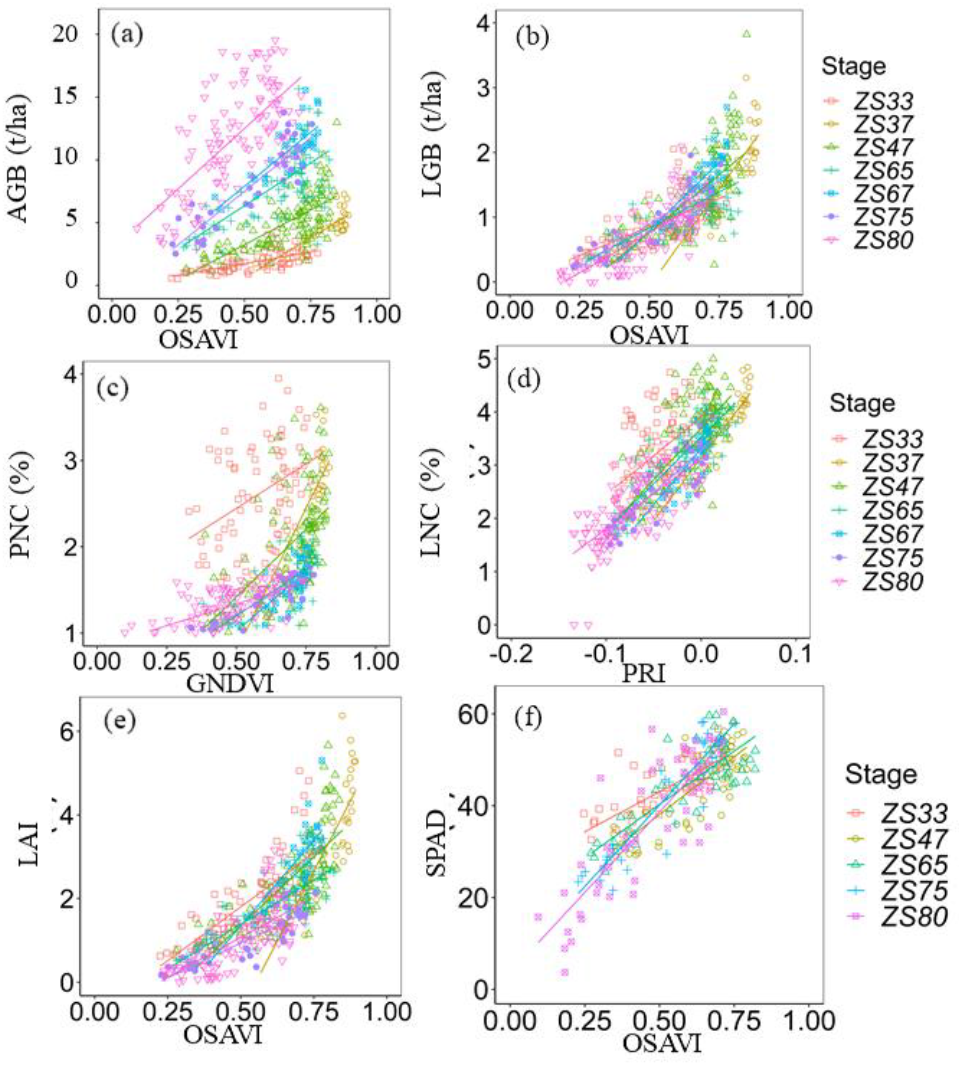
Results of regression analysis between (a) AGB, (b) LGB, (c) PNC, (d) LNC, (e) LAI, and (f) SPAD and VIs at different phenological stages.

**Table 4.** Statistical summary of MLR models applied to the various phenological stages

| ZS | Plant-related parameters |  |  |  |  |  |  |  |  | Leaf-related parameters |  |  |  |  |  |  |  |  |
| --- | --- | --- | --- | --- | --- | --- | --- | --- | --- | --- | --- | --- | --- | --- | --- | --- | --- | --- |
|  | AGB |  |  | PNC |  |  | LGB |  |  | LNC |  |  | LAI |  |  | SPAD |  |  |
|  | k1 | k2 | r | k1 | k2 | r | k1 | k2 | r | k1 | k2 | r | k1 | k2 | r | k1 | k2 | r |
| 33 | 3.68c | 0.38c | 0.61 | 2.06bc | 1.42a | 0.41 | 1.76c | -0.03a | 0.63 | 14.63b | 3.90a | 0.55 | 5.14bc | -0.77b | 0.76 | 34.75c | 24.65a | 0.73 |
| 37 | 7.81c | -1.57d | 0.82 | 6.61a | -2.51e | 0.80 | 5.98a | -3.06c | 0.83 | 23.55a | 3.13d | 0.89 | 13.39a | -7.39f | 0.83 |  |  |  |
| 47 | 9.92bc | -0.41d | 0.72 | 3.13b | -0.14c | 0.58 | 3.56ab | -1.02b | 0.72 | 19.28ab | 3.69b | 0.72 | 6.54bc | -1.92d | 0.75 | 50.26b | 12.94c | 0.64 |
| 65 | 12.12b | 1.89b | 0.64 | 2.14bc | 0.13bc | 0.78 | 2.23b | -0.28bc | 0.73 | 17.51ab | 3.53ab | 0.87 | 4.24c | -0.74a | 0.79 | 46.41bc | 17.06b | 0.83 |
| 67 | 13.84b | 3.27a | 0.78 | 2.76bc | -0.36d | 0.77 | 4.23a | -1.32b | 0.87 | 21.05a | 3.31c | 0.84 | 8.22b | -0.73a | 0.85 |  |  |  |
| 75 | 19.55a | 0.08c | 0.89 | 1.75c | 0.34bc | 0.90 | 2.65b | -0.43ab | 0.88 | 16.80ab | 3.29c | 0.88 | 3.33c | -2.76e | 0.87 | 70.81a | 4.71d | 0.92 |
| 80 | 20.35a | 4.71a | 0.64 | 1.01d | 0.83b | 0.60 | 2.48b | -0.48ab | 0.79 | 13.24b | 3.13d | 0.71 | 3.98c | -0.92c | 0.76 | 70.19a | 3.58e | 0.85 |

Note: A significant level $P < 0.05$ is indicated by a different alphabet. ZS is Zakod's growth stage.

The prediction-model parameters of plant- and leaf-related parameters (Table 4) over various growth stages reveal that the AGB inversion parameters, k_1_ and k_2_, gradually increase with the advancement of the phenological stage, ranging from 3.68 to 20.53 and −1.57 to 4.71, respectively. For PNC models in different growth stages, the parameters k_1_ and k_2_ had opposite trends, ranging from 1.01 to 6.61 and −2.51 to 1.42, respectively. The parameters of AGB and PNC differ significantly over the various growth stages, so the curves of the plant-related parameters as functions of VI in Figure 2 differ significantly over the various growth stages. k_1_ and k_2_ of LGB, LNC, LAI, and SPAD differ over the various growth stages, but, under the combined effect of k_1_ and k_2_, the curves for the leaf-related parameters as functions of VIs are more consistent.

### 3.3 Effect of phenological indicators on predicting plant- and leaf-related parameters

Figure 3 shows the correlation coefficient between the APs of the inversion models for individual growth stages and selected phenological indicators (DAS, DOY, EAT, and ZS). The results shown in Figure 3 indicate that all phenological indicators are strongly correlated parameters (k_1_ and k_2_) of AGB and SPAD inversion models (|r| varies from 0.83 to 0.97). The correlation coefficients |r| of k_1_ for the PNC inversion models with DAS, DOY, GDD, and EAT are 0.61, 0.60, 0.58, and 0.59, respectively, while k_2_ of the PNC inversion models is weakly correlated with the various meteorological factors. Other APs, such as LGB, and LAI, correlate weakly but significantly with the four phenological indicators selected in this paper. Therefore, it is crucial to analyze how the phenological factors affect different estimates of APs.

**Figure 3.**
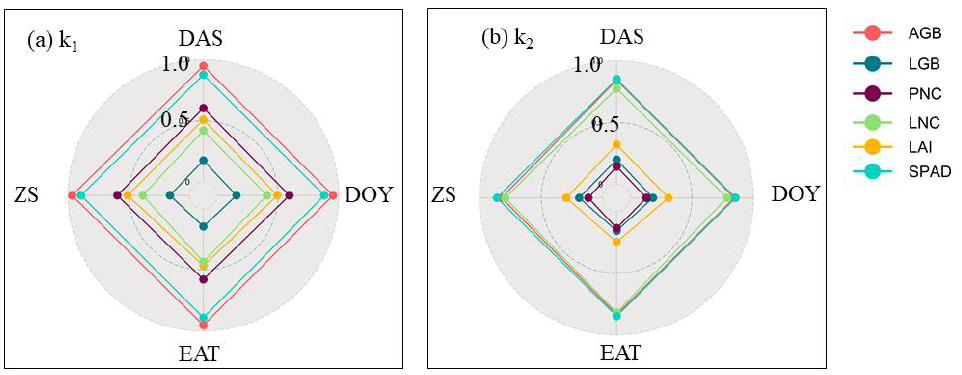
Correlations |r| between k and phenological indicators.

### 3.4 Comparison of agronomic-parameter inversion models with and without phenological indicators as influencing factors

The inversion of AGB and PNC with the phenological indicators considered is more accurate than that obtained with the phenological indicators neglected, but no significant difference appears between the inversion of LGB, LNC, LAI, and SPAD if the phenological indicators are considered (Figure 4. Ignoring the influence of the phenological stage, the accuracy of the AGB model based on VI gives *R*^2^ = 0.02 and RMSE = 4.48 t/ha, which is significantly lower than that obtained with the AGB model combined with phenological information (*R*^2^ = 0.84–0.86 and RMSE = 0.13– 0.50 t/ha). The model based on phenological information has a higher inversion accuracy (*R*^2^ = 0.79–0.87, RMSE = 0.01%–0.07%) than the PNC inversion model based on VI (*R*^2^ = 0.45, RMSE = 0.19%). The range of *R*^2^ for the inversion model of LGB, LNC, LAI, and SPAD is 0.70–0.71, 0.68–0.86, 0.75–0.81, and 0.68–0.70, respectively; and the RMSE is 0.01–0.03 t/ha, 0.02%–0.03%, 0.02–0.07, and 0.47–0.69, respectively.

**Figure 4.**
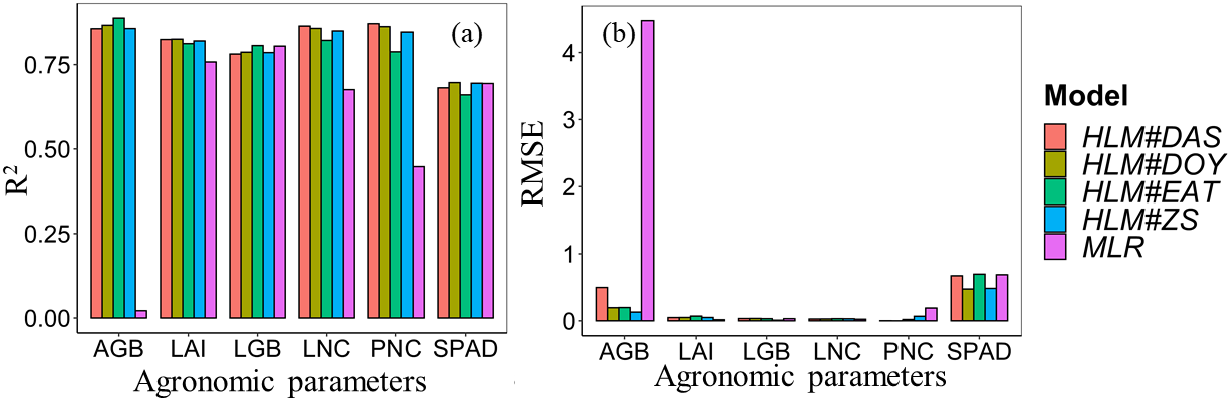
Comparing the accuracy of the inversion model with different agronomic parameters: (a) AGB, (b) LGB, (c) PNC, (d) LNC, (e) LAI, and (f) SPAD.

Figure 5 shows the sensitivity of plant- and leaf-related parameters to VI and phenological stages. The sensitivity concerning plant-related parameters (AGB and PNC) is maximal for phenological indicators (DAS, DOY, EAT, and ZS), whereas the leaf-related parameters (LGB, LNC, LAI, and SPAD) are most sensitive to VIs. The sensitivity index of the phenological information in the AGB and PNC inversion models is 0.56–0.78 and 0.66–0.72, respectively, while the LGB, LNC, LAI, and SPAD inversion models produce sensitivity indexes of 0.01–0.06, 0.01–0.10, 0.02–0.06, and 0.00–0.01, respectively. Optical data can produce leaf information but are limited for detecting plant stems and panicles. With advancing growth stage, the contribution of leaves to canopy information of winter wheat gradually decreases. The information on the growth stage supplements the information not detected by optical data but does not affect the inversion of leaf information.

**Figure 5.**
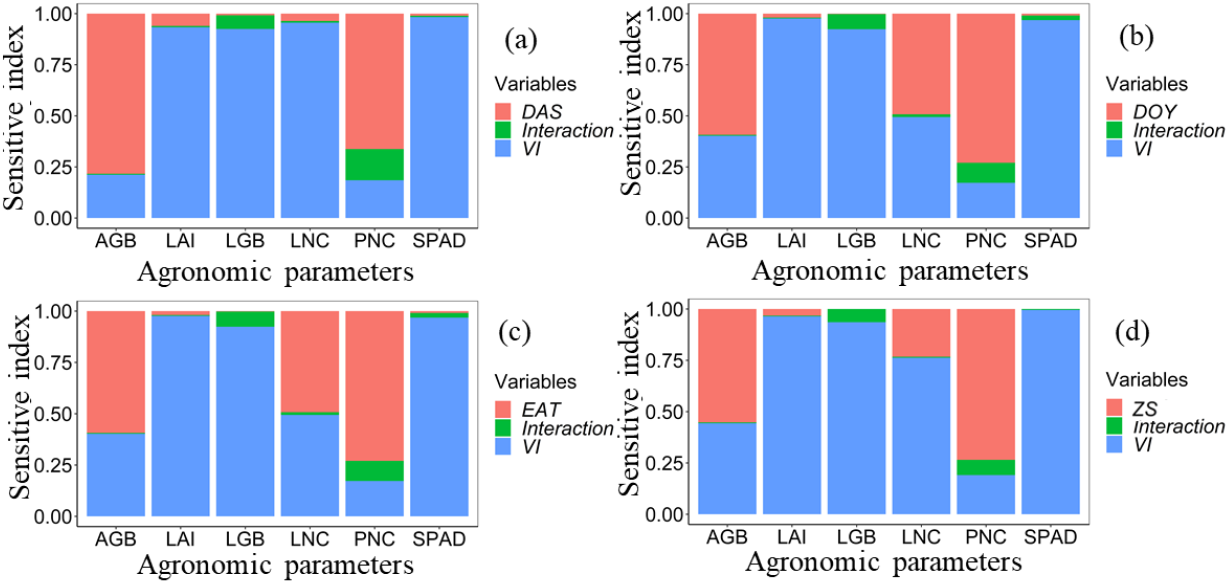
Sensitivity index for AP of HLM

The HLM method to integrate spectral information and the phenological indexes is used to invert the APs. The AGB, LGB, PNC, LNC, LAI, and SPAD models constructed with ZS as independent variable were tested, and the resulting model coefficients appear in Table 5. Figure 6 shows the verification results based on data from the 2019–2020 and 2020–2021 winter wheat growing seasons. Figures 6a and 6c show that, when 7.5 t/ha of AGB and 2.5% of PNC are the thresholds for predicting AGB (RMSE = 3.41 t/ha) and PNC (RMSE = 0.06%) by the MLR model, the result is a clear overestimation (AGB < 7.5 t/ha or PNC <2.5%) and underestimation (AGB > 7.5 t/ha or PNC >2.5%). The HLM model that considers phenological indicators improves upon this overestimation and underestimation when predicting AGB (RMSE = 0.70 t/ha) and PNC (RMSE = 0.01%). The MLR model and HLM model based on LGB, LNC, LAI, and SPAD did not produce significantly dissimilar results for the verification sets.

**Figure 6.**
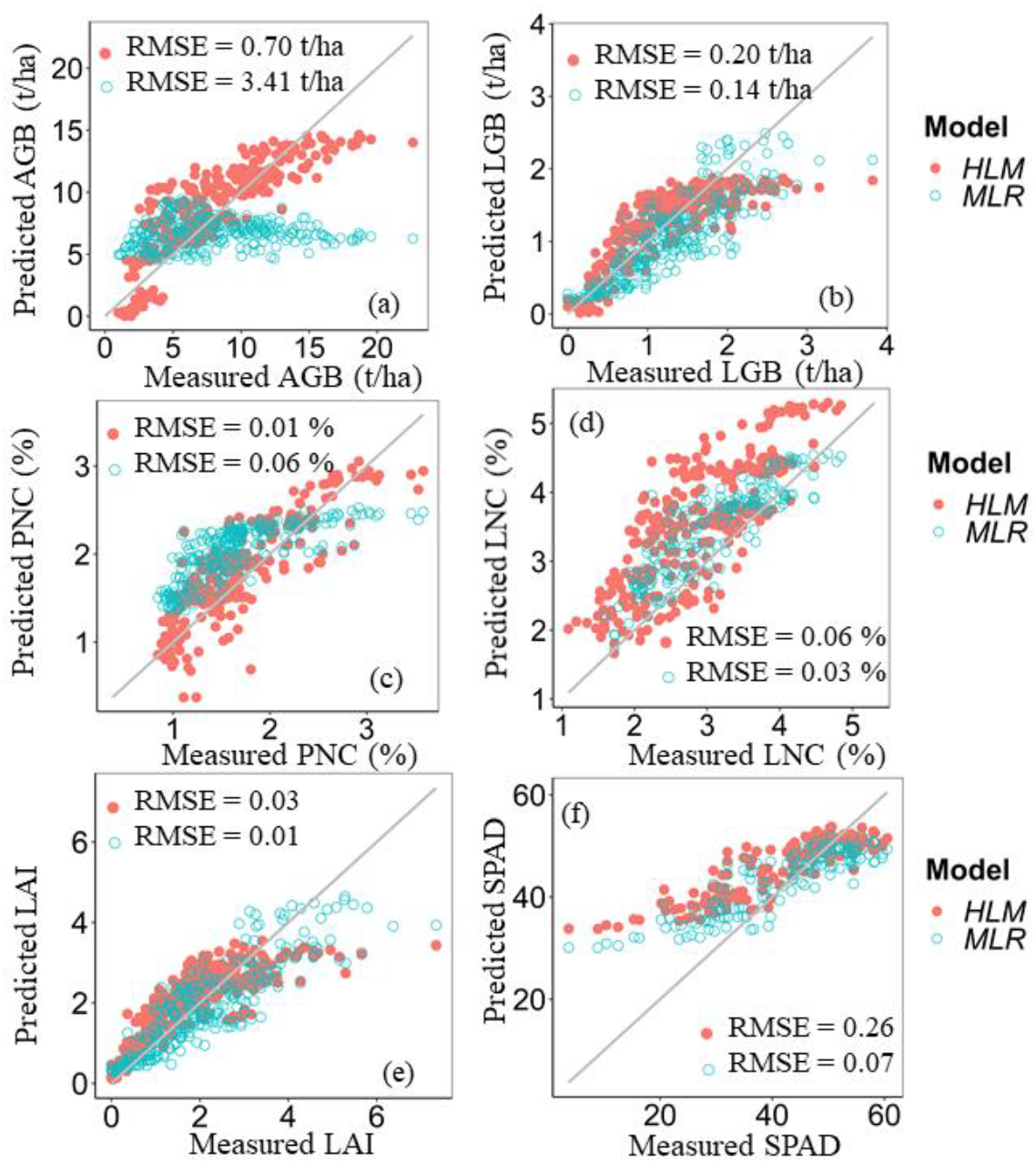
Relationship between measured and predicted (a) AGB, (b) LGB, (c) PNC, (d) LNC, (e) LAI, and (f) SPAD by HLM and MLR methods.

## Discussion

### 4.1 How winter wheat phenological stage affects AP-VI model

Winter wheat is a typical crop with a long growth cycle, large differences in AGB during the different growth stages, and significant nitrogen “dilution” effects, which in turn produce large differences in the time series of AGB and PNC (Justes et al., 1994; Zhao et al., 2020). The leaf-related parameters are like the VI in the phenological stage, which is consistent with previous studies (Zhao et al., 2021; Feng et al., 2019). Figure 7 shows that the phenological stage affects the relationship between leaf-related parameters (LGB or LNC) and plant-related parameters (AGB or PNC). A similar tendency appears when analyzing the correlation between VIs and plant-related parameters (AGB or PNC; see Figs. 2a and 2c). During the growth and development of wheat, its external morphology and internal distribution mechanism undergo a series of significant changes. From the tillering stage to the jointing stage to the flowering stage, the distribution centers of assimilation products are leaves, stems, and spikes (Bidinger et al., 1977; Wagner et al., 1983).

**Figure 7.**
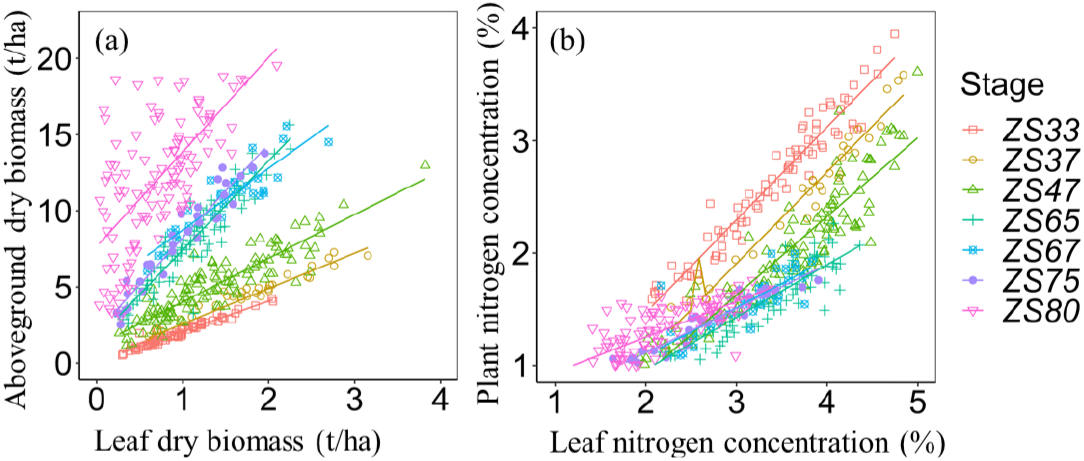
Scatter plots between leaf- and plant-related parameters over different stages: (a) leaf dry biomass and aboveground dry biomass, (b) leaf nitrogen concentration and plant nitrogen concentration.

### 4.2 Response of vegetation indexes to wheat growth stages and agronomic parameters

This study expands efforts to widen the application of VIs for monitoring the APs of winter wheat with multiple phenological stages. The results show that VI inversion of plant-related parameters (AGB and PNC) is more susceptible to the phenological stage than to the leaf-related parameters (LGB, LNC, LAI, and SPAD). In the VI-based AGB and PNC inversion models, k_1_ varies over a large range (from 3.41 to 21.53 and 1.45 to 6.61, respectively). Yue et al. (2019) and Zheng et al. (2018) revealed the phenomenon of plant-related parameters (AGB and PNC) at different growth stages. Classical VIs are constructed mainly to reduce the influence of soil background, aerosol, and light, to reduce other noise, and to limit saturation in the satellite image, whereas less attention is devoted to accurately monitoring phenological effects. Optical data can be used to detect leaves, but their ability to detect plant stems and panicles is limited. As the growth stage advances, the contribution of leaves to canopy information of winter wheat gradually decreases. As shown in Figs. 2 and 7, similar results are obtained when analyzing the correlation of leaf-related indicators and VIs with plant-related indicators at different growth stages. According to the results, the leaf-related APs still perform ideally throughout the phenological stage (Figure 7). Therefore, plant-related parameters at a specific growth stage indirectly give access to leaf-related information. Therefore, the influence of the phenological stage must be understood to construct a VI that overcomes the phenological effect and thereby is able to accurately estimate plant-related parameters.

### 4.3 Should we rely on phenological indicators for parameters inversion?

The results of this study show that the phenological information contributes differently to improving the AP inversion models (Table 5). A comparison of the sensitivity index shows that the phenological information contributes significantly to both plant-related parameters (AGB and PNC; see Figure 5). In a universal model, VIs are not useful for estimating AGB and PNC over the entire growth stage (from tillering to anthesis) because optical data cannot fully detect the vertical structure of the winter wheat canopy (Duan et al., 2019). Four more-accessible indicators, DAS, DOY, EAT, and ZS, are used herein to analyze the contribution of the phenological stage to AP models. The use of the ZS scale as a continuous variable is highly effective for predicting plant-related parameters, regardless of region. Temperature is widely regarded to drive the evolution of the growth stages, but the differences in temperature for crop growth and development differ significantly between regions (Duchem et al., 2008, Zhang and Tao., 2013).

DAS, DOY, and EAT are easily available (Battude et al., 2016; Luo et al., 2020; Manfron et al., 2017) and are suitable for analysis in specific areas, whereas plant-related parameter inversion combined with the ZS stage has a greater potential for use in larger areas. It is thus feasible to use the phenological phase simulation module of the mature crop growth model to predict the regional ZS and then predict the regional AGB, PNC, and even yield. Information such as plant height and texture features that can represent phenological differences at different growth stages was extracted and good results were obtained for inverting plant-related parameters (Yue et al., 2017; Zheng et al., 2018). However, on a regional scale, the current resolution of optical remote sensing restricts the extraction of wheat plant height and texture characteristics.

HLM offers potential advantages in solving the problem of nested data (Li et al., 2019; Hebblewhite and Merrill., 2008), but these advantages do not extend to the inversion of leaf-scale parameters (Figure 6, Table 5). In the HLM models with LGB, LAI, and SPAD, which were constructed by considering phenology, the phenology information exerts a negative effect (i.e., *R*^2^ decreases and RMSE increases). Optical remote sensing mainly provides leaf information, so VIs could lead to acceptable results when analyzing the leaf-related aspects in the multi-phenology stage. At present, all stratifications used to analyze how phenological information affects the inversion of APs are linear models. In the future, different fits, such as nonlinear models or machine learning models, can be analyzed to invert APs over a larger area.

## Conclusion

We report herein on the use of near-ground spectroscopy data in four-year experiments to estimate the plant-related parameters (AGB and PNC) and leaf-related parameters (LGB, LNC, LAI, and SPAD) of winter wheat crops throughout the entire growing season. The results lead to the following main conclusions:

1. Compared with leaf-related parameters (LGB, LNC, LAI, and SPAD), plant-related parameters (AGB and PNC) are more susceptible to the phenological phase in VI-inversion models.
2. The sensitivity index of the phenological information in the AGB and PNC inversion models was 0.56–0.78 and 0.66–0.72, respectively, whereas the sensitivity index for the LGB, LNC, LAI, and SPAD inversion models was 0.01–0.06, 0.01–0.10, 0.02–0.06, and 0.00–0.01, respectively.
3. The AGB inversion model considering ZS (*R*^2^ =0.85, RMSE = 0.13 t/ha; PNC: *R*^2^ = 0.85, RMSE = 0.07%) were better than the inversion models based on VI (AGB: *R*^2^ = 0.02, RMSE = 4.48 t/ha; PNC: *R*^2^ = 0.45, RMSE = 0.19%). No significant difference in accuracy appears between the models based on leaf-related parameters (LGB, LNC, LAI, and SPAD) with phenological information considered and the VI-based inversion models.

## Acknowledgments

This study was supported by the National Key Research and Development Program (2019YFE0125300) and the earmarked fund for China Agriculture Research System (CARS-03).

## Author contributions

Y.Z., Z.L., and G.Y. were involved in study design and research. Y.M., H.F., Y.Z., and S.H. performed the research. Y.Z. analyzed and wrote the paper. The author was responsible for the distribution of materials integral to the findings presented in this article in accordance with the policy described in the Instructions for Authors.

